# Comparison of Techniques for Isolating Small Extracellular Vesicles from *Drosophila* Larval Hemolymph

**DOI:** 10.64898/2026.09.03.748992

**Authors:** Akimi Green, Claudia G. Vásquez, Seong Wook Yang, Young V. Kwon

## Abstract

Extracellular vesicles (EVs) are membrane-delimited nanoparticles, secreted by virtually all tested cell types to mediate intercellular and interorgan communication by transporting biomolecules, such as proteins, nucleic acids, and lipids, to recipient cells. Many EV isolation techniques have been developed for mammalian EVs and exploit specific EV properties, such as size, density, and solubility. Although *Drosophila* has emerged as a simple yet robust animal model for studying fundamental EV biology in its native context, it remains unclear whether commonly used EV isolation techniques can be applied to hemolymph (*Drosophila* blood). In this study, we first provide an in-depth characterization of particles in hemolymph using two complementary particle analysis techniques: dynamic light scattering (DLS) and nanoparticle tracking analysis (NTA). We then evaluate the performance of three commonly used small EV isolation methods—solvent precipitation, ultracentrifugation, and size-exclusion chromatography—for their ability to isolate small EVs from *Drosophila* larval hemolymph. All three methods can enrich small EVs from hemolymph to varying degrees, but none completely remove circulating proteins and lipoproteins. In particular, size-exclusion chromatography yields the purest small EV fractions, as evidenced by the enrichment of *Drosophila* orthologs of human small EV markers, including Tetraspanin 42Ee (Tsp42Ee), Tetraspanin 42Ed (Tsp42Ed), Tetraspanin 96F (Tsp96F), and Annexin B11 (AnxB11), although it produces the lowest protein yield. Altogether, our study provides practical guidance on both selecting an appropriate small EV isolation method and rigorously characterizing isolated small EV fractions according to specific research needs. More broadly, the knowledge gained from this study provides a framework for the isolation and characterization of small EVs in other insects, thereby facilitating future EV research across diverse insect species.

## Introduction

Extracellular vesicles (EVs) are lipid-bilayer delimited nanoparticles released by virtually all cell types across the kingdoms of life (Deatherage & Cookson, 2012; Robinson et al., 2016; Schorey et al., 2015). They were initially discovered in rat reticulocytes (Harding et al., 1983). Subsequent studies in *in vitro* and *in vivo* systems revealed their vital and diverse roles in intercellular and interorgan communications by transporting bioactive cargo, such as ribonucleic acids (RNA), deoxyribonucleic acids (DNA), signaling ligands, enzymes, and metabolites, from donor to recipient cells under various (patho)physiological conditions, including metabolic homeostasis, development, cancer progression, and neurodegeneration (Kalluri & LeBleu, 2020; Kumar et al., 2024; Lin et al., 2023; Payandeh et al., 2024; Puthiyattil et al., 2026; Salomon et al., 2022; Sheng et al., 2025; Yáñez-Mó et al., 2015; Yates et al., 2022).

EVs are highly heterogeneous in nature, ranging from 10 to 20,000 nm in diameter, carrying diverse cargo, and originating from distinct subcellular compartments (Carney et al., 2025; Kalluri & LeBleu, 2020; Silva et al., 2025; van de Wakker et al., 2023; Xu et al., 2025). While EV classification is continuously evolving, EVs are generally categorized into three distinct groups based on size and biogenesis mechanism: exosomes, ectosomes, and apoptotic bodies (Chen & Yang, 2024) (Figure 1A). Exosomes are endosome-derived EVs that range in diameter from 30 to 150 nm (Bebelman et al., 2018; Borges et al., 2013; Raposo & Stoorvogel, 2013; Zaborowski et al., 2015). Exosome biogenesis starts with maturation of early endosomes into late sorting endosomes, which subsequently invaginate to form multivesicular bodies (MVBs). Fusion of the MVB membrane to the plasma membrane results in the release of exosomes into the extracellular milieu (Van Niel et al., 2018). Ectosomes, also called microvesicles, are plasma membrane-derived vesicles, ranging in size from 50 nm to 1 µm (Jin et al., 2022; Singh et al., 2025; Tricarico et al., 2017). Ectosome shedding from the plasma membrane requires actomyosin dependent fission and release (Tricarico et al., 2017). Apoptotic bodies are large (0.5–5 µm) plasma membrane-derived vesicles released from dying cells in the last stage of apoptosis (Yu et al., 2023).

**Figure 1.**
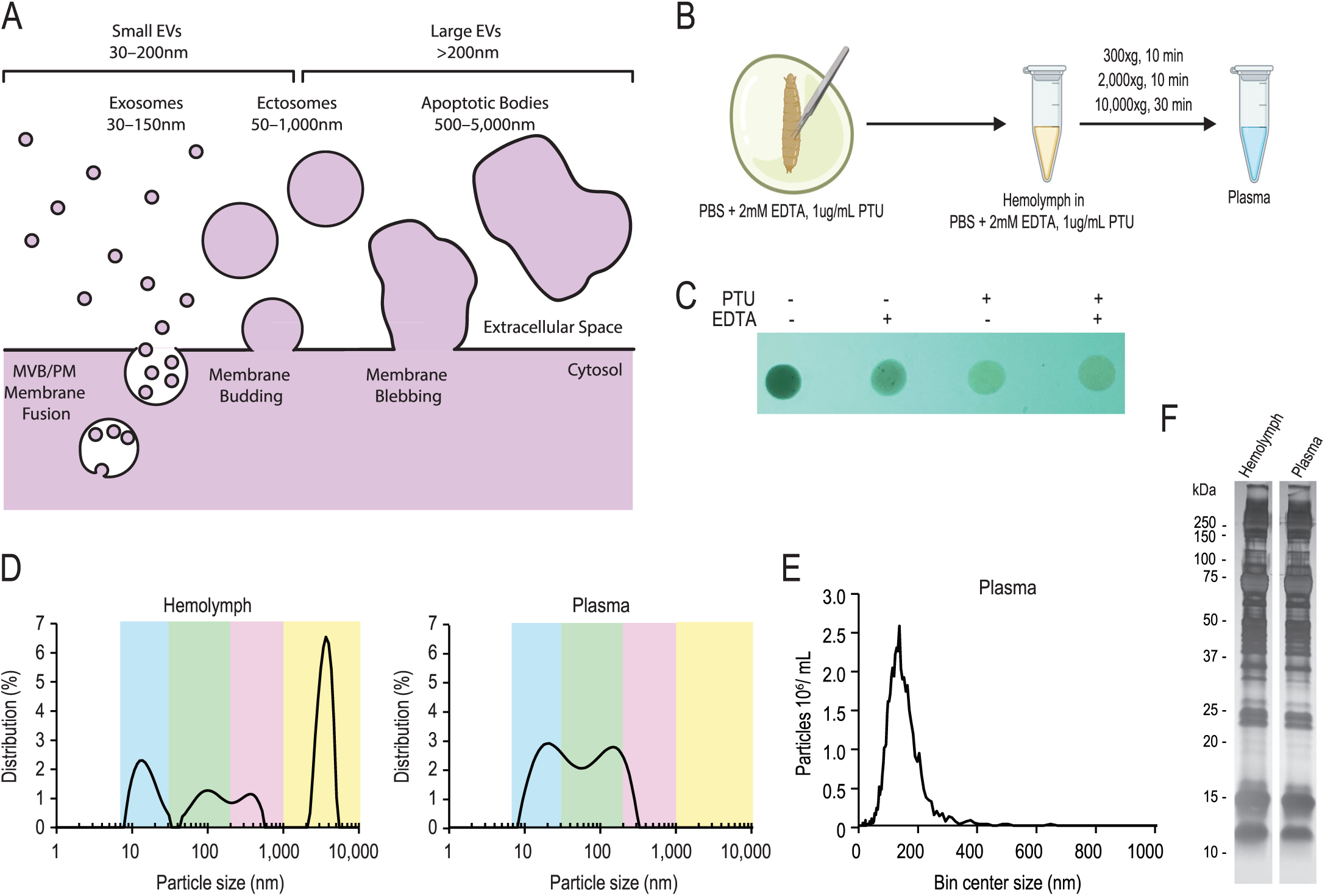
Preparing *Drosophila* hemolymph for enrichment of small EVs. (**A**) Illustration of exosomes, ectosomes, and apoptotic bodies. Small and large EVs are defined based on their size. (**B**) Schematic of hemolymph collection and plasma preparation for small EV isolation. (**C**) Comparison of hemolymph melanization. 2 mM EDTA and 1 µg/ml phenylthiourea were added to phosphate buffered saline. (**D**) Size distribution of particles in hemolymph (left) and plasma (right). Particle size distribution was determined by DLS using three manual replicate measurements, with each run lasting 30 s. (**E**) NTA of plasma. (**F**) Silver staining of hemolymph and plasma.

While the biogenesis mechanisms of exosomes, ectosomes, and apoptotic bodies are defined, analysis of circulating EVs does not necessarily involve interrogation of EV biogenesis mechanisms. After secretion into the extracellular milieu, it is challenging to identify whether the EV population of interest originated from the endosomal system or the plasma membrane. Thus, circulating EVs can be divided into 2 subgroups: small EVs and large EVs. Small EVs refer to EVs 30–200 nm in diameter, which predominantly include exosomes and small ectosomes, whereas large EVs refer to EVs >200 nm, such as large ectosomes and apoptotic bodies (Welsh et al., 2024). Since we do not investigate EV biogenesis mechanisms in this study, we chose to abide by this size-based classification (Figure 1A).

Due to the complexity of *in vivo* mammalian systems, studies elucidating the function of mammalian EVs have largely been done in tissue culture systems. Investigation of the biological function of EVs in non-mammalian species has great potential to elucidate both conserved and uniquely adapted roles of EVs in an animal (Liegertová & Janoušková, 2023). The powerful genetic toolkit and conserved pathways of *Drosophila melanogaster*, make it an ideal model for investigating fundamental EV biology *in vivo* (Cotterill & Yamaguchi, 2023; Lloyd & Taylor, 2010; Reiter et al., 2001).

EVs have been isolated from *Drosophila* cell culture supernatant and characterized via proteomics and transcriptomics (Beckett et al., 2013; Koppen et al., 2011; Lefebvre et al., 2016). These studies identified similarities in size, morphology, and cargo content between *Drosophila* and human EVs (Beckett et al., 2013; Koppen et al., 2011; Lefebvre et al., 2016). Additionally, several studies have highlighted the physiological function of EVs in *Drosophila* (Blanchette et al., 2022; Camelo et al., 2022; Gross et al., 2012; Hurbain et al., 2022; Koles et al., 2012; Lee et al., 2026; Linnemannstöns et al., 2022). Live imaging of the developing *Drosophila* embryo showed that small GTPases Arl3, Rab27, and Rab35 are required for the formation and release of EVs into the tracheal tube lumen (Camelo et al., 2022). Further, a subpopulation of these EVs were shown to be spatially and temporally associated with tracheal tube fusion events. Inhibition of EV secretion resulted in dorsal branch fusion defects (Camelo et al., 2022). The larval neuromuscular junction (NMJ) is another robust *in vivo* model for investigating synaptic EV biology (Koles et al., 2012). The presynaptic terminal releases endosome-derived EVs into the synaptic space in a manner dependent on Syntaxin 1A, a plasma membrane SNARE protein, and the recycling endosome machinery components Myo5 and Rab11 (Koles et al., 2012). An additional study revealed that endocytic machinery is required for the trafficking and sorting of MVB cargo at the NMJ, with mutations to the endocytic machinery resulting in synaptic plasticity defects (Blanchette et al., 2022). In the larval wing disc, EVs contain and transport morphogens, such as Wnt and Hedgehog. Loss of Prominin-like protein (PromL) results in abnormal microvilli and inhibits EV secretion, and consequently, impairs long-range Hedgehog signaling (Gross et al., 2012; Hurbain et al., 2022). A recent study demonstrated that the biogenesis of large EVs from malignant cells is a conserved process in *Drosophila* (Lee et al., 2026). The tumor cell cytosolic DNA response, mediated by the cyclic GMP-AMP synthase (cGAS)-stimulator of interferon genes (STING) pathway, drives large EV biogenesis in a JNK and FAK-dependent manner (Lee et al., 2026).

Although these studies underscore the prevalence and importance of EVs in *Drosophila* development and disease, the isolation of small EVs from biofluids, such as *Drosophila* ‘blood’ known as hemolymph, for biochemical and functional analyses remains challenging. In this study, we sought to define the benefits and limitations of widely used small EV isolation methods when applied to larval *Drosophila* hemolymph. We performed thorough characterization of small EV fractions prepared via solvent precipitation, ultracentrifugation, and size-exclusion chromatography. Our study provides a rigorous assessment of the performance of these small EV isolation methods, offering guidance on selecting an appropriate method for specific research needs while ensuring accurate analysis of the size and concentration of isolated small EVs.

## Results

### Optimization of hemolymph collection buffer to minimize coagulation and melanization

Hemolymph coagulation and melanization are two of the most immediate physical defense mechanisms triggered upon wounding or infection (Theopold et al., 2004). Injury induces a rapid coagulation of hemolymph, forming a soft clot to plug the epithelial breach and to trap circulating pathogens (Theopold et al., 2004). Cross-linking of clotting factors, such as Hemolectin, Fondue, and lipoproteins, is primarily catalyzed by transglutaminases (Dushay, 2009). Melanization is the deposition of melanin at the site of infection or injury, playing a role in wound healing, hardening the clot, and encapsulating larger parasites (Bidla et al., 2005; Sheehan et al., 2018). Phenoloxidase (PO) is a key enzyme in the melanin synthesis cascade (Bidla et al., 2005).

Previous studies showed that clots can trap EVs and negatively impact the performance of small EV isolation procedures (Arakelyan et al., 2016; Zhang et al., 2022). Thus, anticoagulants, such as ethylenediaminetetraacetic acid (EDTA), are widely used during blood collection for EV isolation (Aatonen et al., 2014; Arakelyan et al., 2016). In *Drosophila*, EDTA is also used to prevent coagulation by chelating Ca^2+^, which is essential for the activity of transglutaminases (Ahvazi et al., 2003; Hand et al., 1985). Melanization can be reduced by supplementing the hemolymph collection buffer with phenylthiourea (PTU), an inhibitor of PO (Ryazanova et al., 2012). We collected hemolymph by carefully puncturing third-instar larvae with forceps and letting them bleed into filtered PBS (Figure 1B). After a 30-minute incubation at room temperature, collected hemolymph darkened noticeably, indicating that coagulation and melanization had occurred (Figure 1C). Supplementing PBS with 2 mM EDTA, 1µg/ml PTU, or both significantly reduced hemolymph darkening (Figure 1C). Therefore, we decided to use PBS supplemented with both 2 mM EDTA and 1µg/ml PTU as the hemolymph collection buffer. To further minimize melanization and coagulation, hemolymph collection and all subsequent procedures were performed on ice or at 4 °C.

### Low speed centrifugation removes large particles and cell debris from *Drosophila* hemolymph, preparing for small EV isolation

*Drosophila* hemolymph is a complex biofluid composed of multiple circulating components, including proteins, lipoproteins, and hemocytes (*Drosophila* macrophage-like cells) (Handke et al., 2013). To obtain a comprehensive assessment of particle sizes in hemolymph, we analyzed hemolymph samples collected from wild-type larvae using dynamic light scattering (DLS), a technique commonly used to estimate particle size by measuring light scattered from particles in solution. While DLS can detect particles ranging from approximately 0.3 nm to 10 µm in diameter, DLS measurements are biased toward larger particles since larger particles scatter more light than smaller particles (Kim et al., 2019). Therefore, DLS profiles provide an overall particle size distribution but do not accurately reflect the relative abundance of particles of different sizes.

Interestingly, our DLS analysis detected 4 distinct particle populations, with peaks around 15, 100, 400, and 3,500 nm (Figure 1D, left). To facilitate visualization of different particle populations, we color-coded the background of the DLS graphs throughout the manuscript: blue for 7–30 nm, green for 30–200 nm, pink for 200–1,000 nm, and yellow for 1,000–10,000 nm (Figure 1D). Lipophorins are functionally analogous to mammalian lipoproteins. In some insects, such as silkworm, cockroach, and locust, lipophorins are measured to be 13–16 nm in diameter (Chino, 1982). Thus, the particles measured at approximately 15 nm (blue) are likely to be lipophorins (Figure 1D, left). Mammalian exosomes typically range from 30 to 150 nm in diameter. The mean diameter of exosome-like EVs isolated from *Drosophila* S2 cells was approximately 100 nm (Beckett et al., 2013; Kerr et al., 2018; Lee et al., 2023). Thus, the particle population that peaks at approximately 100 nm (green) is reminiscent of exosomes (Figure 1D, left). Microvesicles are large EVs ranging from 100 nm to 1,000 nm. Chylomicrons are large and low-density lipoproteins found in the circulatory system ranging in size from 75 nm to over 1,200 nm. Thus, the particle population that peaks at approximately 400 nm (pink) may correspond to microvesicles, chylomicron-like lipoproteins or both (Figure 1D, left). Based on their size distribution, the very large particles ranging from 2,000 nm to 6,000 nm (yellow) may represent cell debris and small cells (Figure 1D, left).

Our DLS analysis showed that hemolymph contains cell debris and particles larger than conventional small EVs (Figure 1D, left). Preclearing samples to remove large particles is a routine step in exosome isolation (Dilsiz, 2024; Konoshenko et al., 2018; Sidhom et al., 2020a). Therefore, we tested whether a series of low-speed centrifugation steps could remove large particles by pelleting them (Figure 1B). Three sequential centrifugations at 300 × *g* for 10 min, 2,000 × *g* for 10 min, and 10,000 × *g* for 30 min efficiently removed particles larger than 300 nm, resulting in two major particle populations (Figure 1D, right). Unlike microvesicles and cell debris, chylomicron-like lipoproteins have very low density and cannot be pelleted by low-speed centrifugation. Because our low-speed centrifugation procedure efficiently removed particles with a peak size of ∼400 nm (Figure 1D, left), large lipoproteins are unlikely to account for particles in this size range. Instead, these particles are likely to represent large EVs, such as microvesicles, and/or small cell debris. We refer to the resulting large particle-depleted fraction as ‘plasma’ and used it for all downstream analyses.

NTA combines laser light scattering microscopy with a charge-coupled device camera, allowing real-time visualization of individual particles dispersed in a sample (Filipe et al., 2010). Since the volume of the sample chamber is known, NTA enables determination of particle size as well as concentration. NTA analysis showed that the majority of particles in the plasma ranged from 50 to 250 nm (Figure 1E), indicating that the particle population peak at ∼400 nm (Figure 1D, pink) was efficiently removed. It should be noted that NTA cannot reliably measure particles smaller than 30 nm because of its lower detection limit (Filipe et al., 2010; Lu & Murphy, 2018). Thus, smaller particles, such as lipoproteins, may be undetected when measured by NTA alone. When we analyzed hemolymph and plasma by SDS-PAGE followed by silver staining, their overall protein profiles were similar (Figure 1F), suggesting that the large particles (Figure 1D, pink and yellow) constitute only a small fraction of the hemolymph.

### Solvent precipitation enriches for both small and large EVs

Solvent precipitation (SP) using commercially available kits is a common method used to isolate small EVs. SP enriches EVs by adding an organic solvent to alter EV solubility, causing EV aggregation. EVs are subsequently pelleted by low-speed centrifugation. We treated *Drosophila* plasma using a commercial isolation kit, resuspended the final pellet in PBS, and referred to this resuspended pellet as ‘SP sEV’ (Figure 2A).

**Figure 2.**
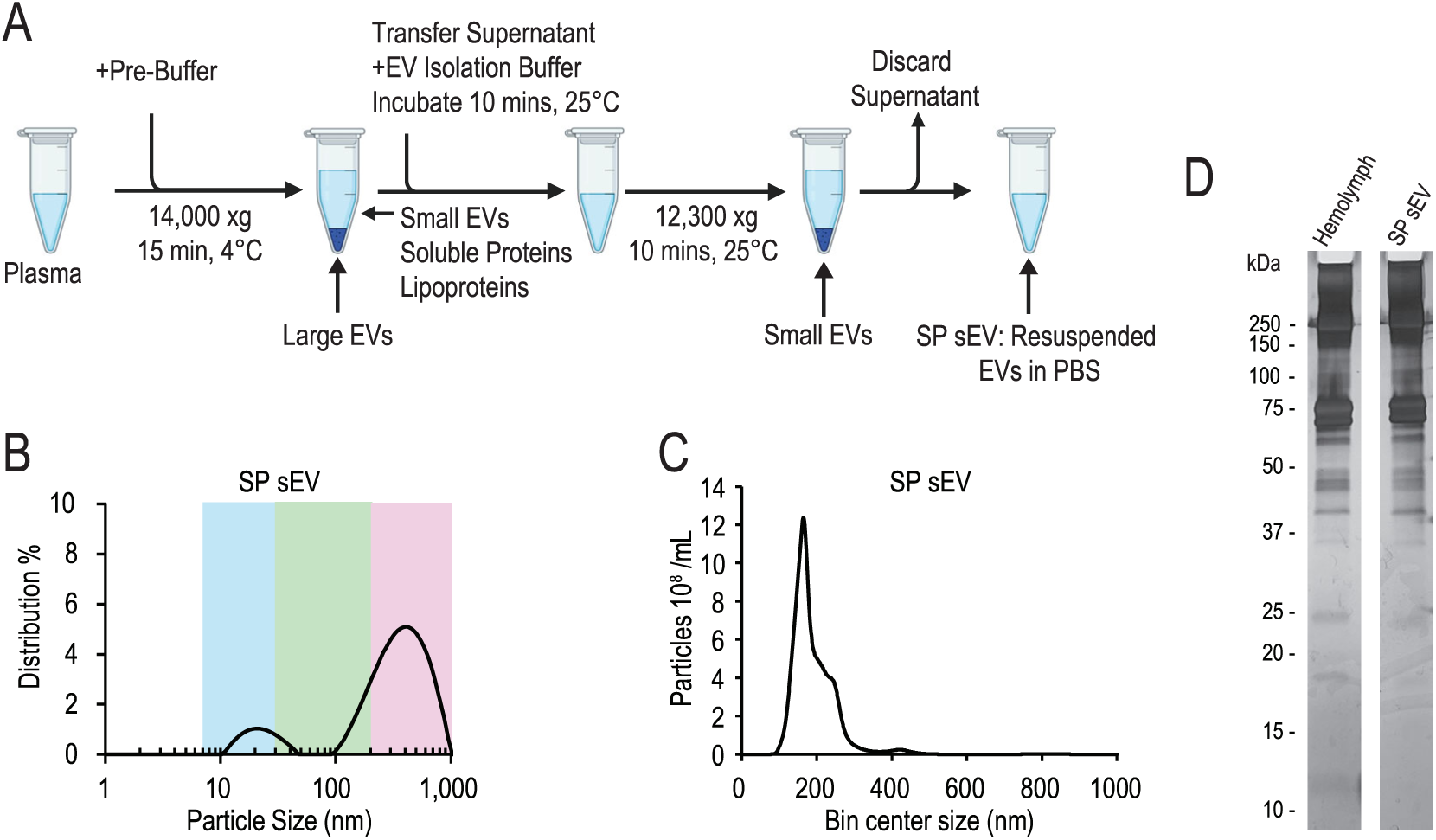
Solvent precipitation enriches a broad range of particles. (**A**) Schematic of *Drosophila* circulating small EV isolation by solvent precipitation. (**B**) Intensity-weighted size distribution of the SP small EV (SP sEV) fraction measured by DLS on manual repetition series, 30s per run for 3 runs. (**C**) Particle size distribution of SP sEV measured by NTA. (**D**) Silver staining of hemolymph and the SP sEV fraction. Proteins were separated by SDS-PAGE and visualized by silver staining.

To assess how efficiently the SP procedure enriched small EVs, we analyzed the SP sEV fraction using DLS. While we detected a distinct particle population peak at approximately 30 nm in the SP sEV fraction (Figure 2B, blue), the dominant population of particles was between 200–600 nm (Figure 2B, pink). Since the small particles (< 30 nm) are much smaller than small EVs, they scatter significantly less light. Thus, their presence might be underestimated in our DLS analysis. Our DLS analysis detected few particles within the defined small EV size range of 30–200 nm (Figure 2B, green). Notably, NTA analysis showed that the majority of detectable particles in the SP sEV fraction ranged from 100 nm to 300 nm, with a peak at approximately 170 nm (Figure 2C).

If our SP procedure successfully isolated small EVs, small EV-specific proteins should be enriched in the SP sEV fraction relative to the hemolymph. Moreover, the SP sEV fraction should contain fewer circulating proteins. However, when we compared the protein profiles of hemolymph and the SP sEV fraction by silver staining, we observed a striking similarity (Figure 2D). With the exception of several proteins in the 10–25 kDa range that were more abundant in hemolymph, the protein profiles were nearly identical. Given the similarity between the hemolymph and the SP sEV fraction, we speculate that the SP procedure co-precipitates abundant circulating proteins.

Our data demonstrate that SP enriches small EVs; however, DLS and NTA results show that the SP sEV fraction also includes particles larger than conventional small EVs (Figure 2B and C). This may indicate that the SP procedure also efficiently enriches large EVs. Alternatively, the SP procedure might alter the properties of small EVs by promoting their fusion/aggregation, enhancing their interactions with circulating proteins, and/or causing them to form complexes with the EV precipitation reagent (Lobb et al., 2015; Van Deun et al., 2014). The SP procedure was unable to completely remove a population of small particles (< 30 nm). Moreover, our protein profile comparison suggests that the SP procedure may be unable to remove circulating proteins.

### Ultracentrifugation enriches small EVs and lipoprotein-like particles from hemolymph

Ultracentrifugation (UC) is another common small EV enrichment method, which can separate small EVs from soluble proteins and lipoproteins based on their density differences (Brennan et al., 2020; Clos-Sansalvador et al., 2022). To enrich small EVs from hemolymph, we adapted a conventional UC protocol for isolating exosomes in cell culture media and biological fluids (Théry et al., 2006) (Figure 3A). We spun plasma at 100,000 × *g* for 70 minutes to precipitate small EVs and remove soluble proteins and low-density particles, such as lipoproteins. We removed the supernatant (SN1) and resuspended the resulting pellet in filtered PBS, followed by a second centrifugation at 100,000 × *g* for 70 minutes to remove residual soluble proteins and low-density particles. After removing the supernatant of the second centrifugation (SN2), we resuspended the pellet in filtered PBS. We referred to this resuspended pellet as ‘UC sEV’ (Figure 3A).

**Figure 3.**
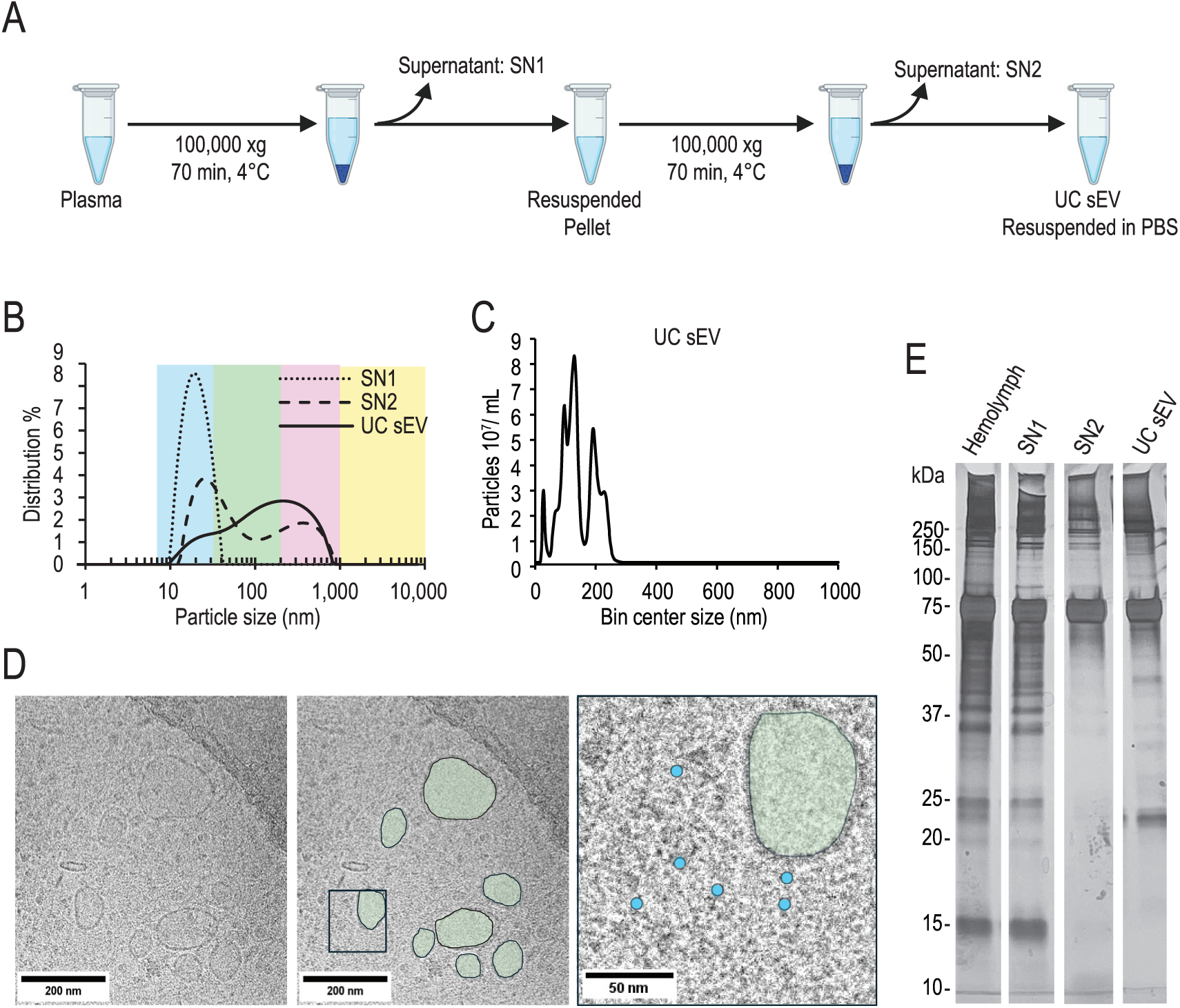
Ultracentrifugation enriches small EV-sized particles. (**A**) The ultracentrifugation procedure for enriching small EVs in *Drosophila* plasma. SN1: supernatant after the first 100,000 x g centrifugation; SN2: supernatant after the second 100,000 x g centrifugation; UC sEV: pellet collected after two 100,000 x g centrifugation steps and resuspended in filtered PBS. (**B**) Particle size distribution of SN1, SN2, and UC sEV fractions. Data was collected by DLS using three manual replicate measurements, with each run lasting 30 s. (**C**) NTA of the UC sEV fraction. (**D**) Representative cryo-EM image of the UC sEV fraction. Multiple membrane-bound structures ranging from 40–180 nm in diameter are observed and are highlighted in green on the right insets. Particles smaller than sEVs are highlighted in blue in the right inset. Scale bar: 200 nm. (**E**) Proteins from each fraction were separated by SDS-PAGE and visualized by silver stained. Arrowheads indicate proteins enriched in the UC sEV fraction compared with the hemolymph, SN1, and SN2 fractions.

To assess which sized particles are enriched and lost throughout the UC procedure, we compared DLS analyses of the SN1, SN2, and UC sEV fractions (Figure 3B). We detected a distinct particle population with a peak at approximately 20 nm in SN1 (Figure 3B, SN1, blue) reminiscent of the small particles observed in the hemolymph and plasma samples (Figure 1D, blue), indicating that these particles were of low-density. In addition to their size distribution, the sedimentation properties of these particles are reminiscent of lipoproteins (Brennan et al., 2020). In SN2, particles smaller than 30 nm were still detected (Figure 3B, SN2, blue), and we also detected particles ranging from 30 nm to ∼1,000 nm (Figure 3B, SN2, green and pink). Analysis of the UC sEV fraction showed presence of particles smaller than 30 nm (Figure 3B, UC sEV, blue) and particles spanning 30–1,000 nm (Figure 3B, UC sEV, green and pink). While DLS analysis showed the presence of particles larger than 300 nm in the UC sEV fraction (Figure 3B, UC sEV), these particles were not detected by NTA, suggesting that their concentration was negligible (Figure 3C). The particles in the UC sEV fraction ranged from approximately 50 nm to 250 nm, with two distinct peaks at approximately 130 nm and 200 nm (Figure 3C). These results suggest that the UC procedure enriches small EV-sized particles; however, it does not completely remove particles smaller than 30 nm.

To determine if the particles in the UC sEV fraction are small EVs, we performed cryo-electron microscopy on this fraction. We detected lipid bilayer-delimited particles measuring 30–200 nm in diameter, providing evidence that this fraction contains small EVs (Figure 3D, green). We also detected an abundance of 10–20 nm particles (Figure 3D, blue). This result supports our DLS data, which indicated the presence of particles smaller than 30 nm in the UC sEV fraction.

The protein profile of the SN1 faction was almost identical to that of hemolymph, suggesting that most soluble proteins in hemolymph remained in the supernatant after the first round of UC (Figure 3E). Most proteins smaller than 75 kDa in the hemolymph and the SN1 fraction were significantly reduced in the SN2 fraction (Figure 3E). These proteins might constitute soluble circulating proteins that were not sedimented by UC. Considering that our DLS analysis detected a particle population with a peak at approximately 30 nm, the protein profile may reflect the enrichment of these particles. The main protein components of *Drosophila* lipophorins, apolipophorin I (ApoLI) and apolipophorin II (ApoLII) are approximately 250 and 75 kDa, respectively (Ryan & Van Der Horst, 2000; Zhao et al., 2020). We observed ∼250 and ∼75 kDa proteins in the SN1 and SN2 fractions (Figure 3E), further supporting that these particles smaller than 30 nm were lipophorins. We noticed that these 75 kDa proteins were also main constituents of the UC sEV fraction, consistent with our cryo-EM observations (Figure 3D and E).

Altogether, our data show that UC can be used to enrich circulating small EVs from hemolymph. Two consecutive rounds of UC successfully removed a large quantity of soluble circulating proteins and lipophorins from hemolymph. Nevertheless, a substantial quantity of lipophorins remained in the final pellet. The inability to separate small EVs from lipophorins may be due to their similarity in density (Yuana et al., 2014). Alternatively, lipophorins may co-sediment with small EVs during UC because they form a complex with small EVs, as previously reported in humans (Ghebosu et al., 2024; Tóth et al., 2021). In conclusion, while UC enriches small EVs from *Drosophila* plasma, additional purification steps are required to further remove lipophorins.

### Genetic depletion of fat body-derived Apolipophorin reduces lipophorins in UC sEV fraction

Our particle size analysis has shown the presence of lipophorins in both the SP and UC sEV fractions, suggesting that these conventional techniques cannot completely separate lipophorins from small EVs. A previous study showed that Apolipophorin (Apolpp) depletion in the fat body significantly reduced the presence of Apolpp in hemolymph (Palm et al., 2012). Thus, we assessed whether genetically depleting Apolpp in the fat body can reduce lipophorins in the small EV fraction prepared by UC.

Apolpp is synthesized in the *Drosophila* fat body and is post-translationally cleaved into ApoLI and ApoLII (Brankatschk & Eaton, 2010). Expression of *apolpp RNAi* in the fat body, using *Cg-GAL4* (*Cg>apolpp RNAi*), reduced ApoLII levels in the hemolymph (Figure 4A). Notably, we observed a consistent reduction of ApoLII in all four fractions, including the UC sEV fraction, when Apolpp was depleted in the fat body (Figure 4A). Although Apolpp depletion in the fat body did not noticeably alter the overall particle size distribution, the small EV population observed in both the hemolymph and UC sEV fraction became more prominent after Apolpp depletion relative to the lipophorin populations (Figure 4B and C). Notably, NTA revealed that the depletion of Apolpp in the fat body increased the small EV yield by approximately four-fold (Figure 4D, left). The average particle size remained consistent (∼150 nm) regardless of Apolpp depletion (Figure 4D, right). The protein profile of the UC sEV fractions also appeared different between the control and *apolpp RNAi* conditions (Figure 4E). One of the cleavage products of Apolpp, ApoLII, runs at approximately 75 kDa (Ryan & Van Der Horst, 2000; Zhao et al., 2020). Thus, the significant reduction of the protein bands at ∼75 kDa in the *Cg*>*apolpp RNAi* condition may be the result of Apolpp knockdown (Figure 4E, arrowheads and brackets). Further, there is a noticeable enrichment of proteins at 10–40 kDa in the *apolpp RNAi* UC sEV fraction compared to the control fraction (Figure 4E). Altogether, our data show that Apolpp depletion in the fat body reduces lipophorins in the hemolymph and UC sEV fractions, suggesting that combining genetic depletion of Apolpp with conventional small EV isolation techniques may be a way to increase the purity of small EV fractions.

**Figure 4.**
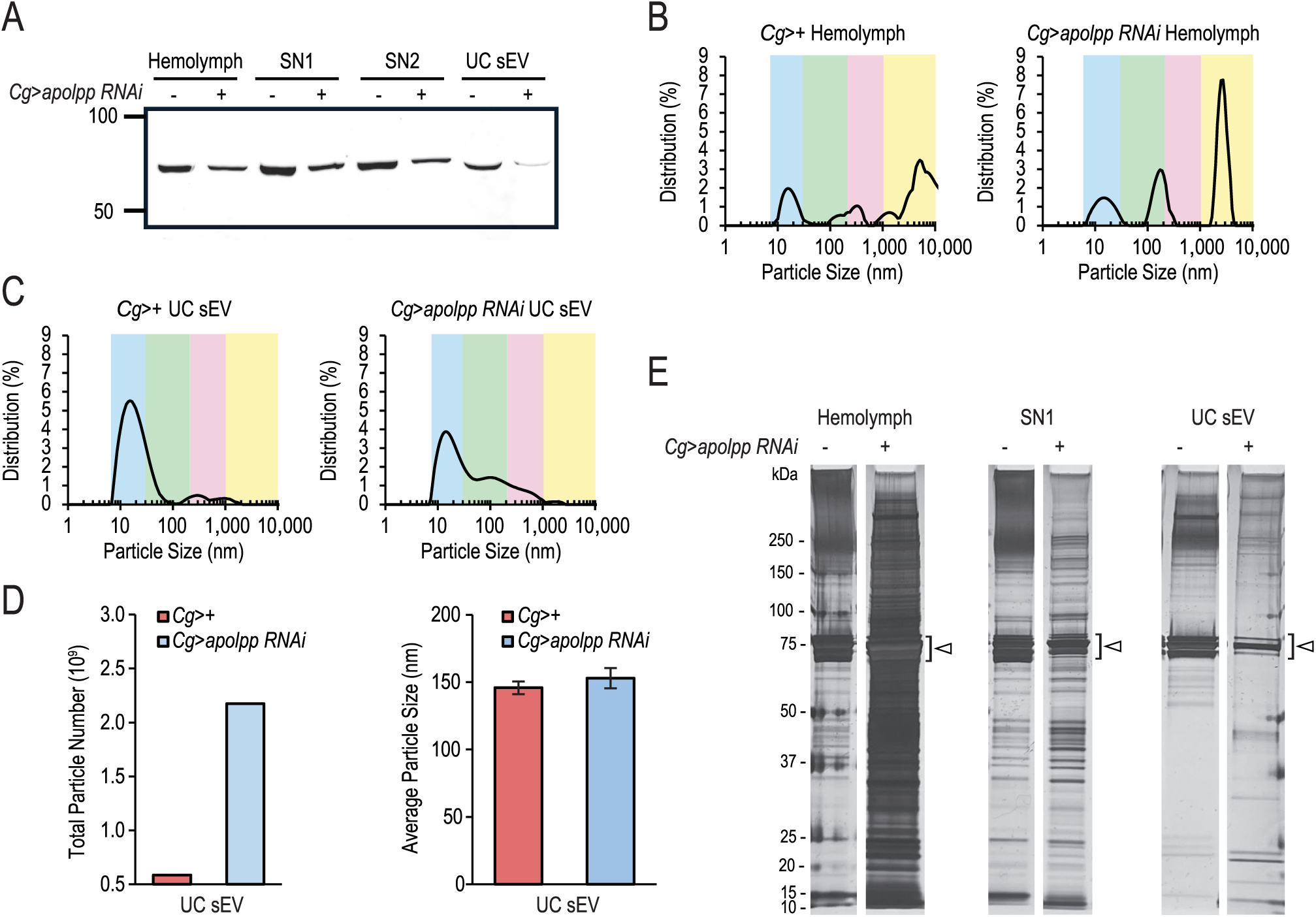
Genetic depletion of Apolpp in the fat body reduces apolipoproteins in hemolymph and the UC sEV fraction. (**A**) Immunoblotting for ApoLII in UC fractions. The expression of apolpp RNAi in the fat body decreased ApoLII in all four UC fractions. *Cg-GAL4* was used to drive *apolpp RNAi* in the fat body. (**B-C**) DLS analysis of hemolymph (**B**) and UC sEV fractions (**C**) from *Cg>+* (control) and *Cg>apolpp RNAi* larvae. Data was collected by DLS using three manual replicate measurements, with each run lasting 30 s. (**D**) Total particle number and particle size in hemolymph and the UC sEV fractions. Data was collected by NTA from three videos. Particle sizes are shown as mean ± SEM. (**E**) Silver staining of hemolymph, SN1, and UC sEV fractions collected from *Cg>+* and *Cg>apolpp RNAi* larvae.

### Size exclusion chromatography enriches small EV-sized particles

Size exclusion chromatography (SEC) is also commonly used for separating small EVs in cell culture media and blood samples (Sidhom et al., 2020b). To determine whether SEC can also separate small EVs in hemolymph, we processed *Drosophila* plasma using a commercial small EV isolation column (Figure 5A). Full fractionation of plasma occurred over 25 fractions (Supplementary Figure 1A). We used DLS to define the particle populations of each fraction. Large particles with a peak ∼1000 nm were eluted in fraction 5 and small EV-sized particles were detected in fractions 6–9 (Figure 5B, Supplementary Figure 1A). Lipophorin-like small particles (10–30 nm) started eluting in fraction 10, and small EV-sized particles became almost absent in fraction 13 (Supplementary Figure 1A). Based on these results, we decided to further analyze fractions 7–9 with NTA (Figure 5C). Consistent with our DLS data, most particles in all three fractions were between 40–150 nm (Figure 5C). Fraction 8 contains the highest concentration of particles compared to fractions 7 and 9 (Figure 5C).

**Figure 5.**
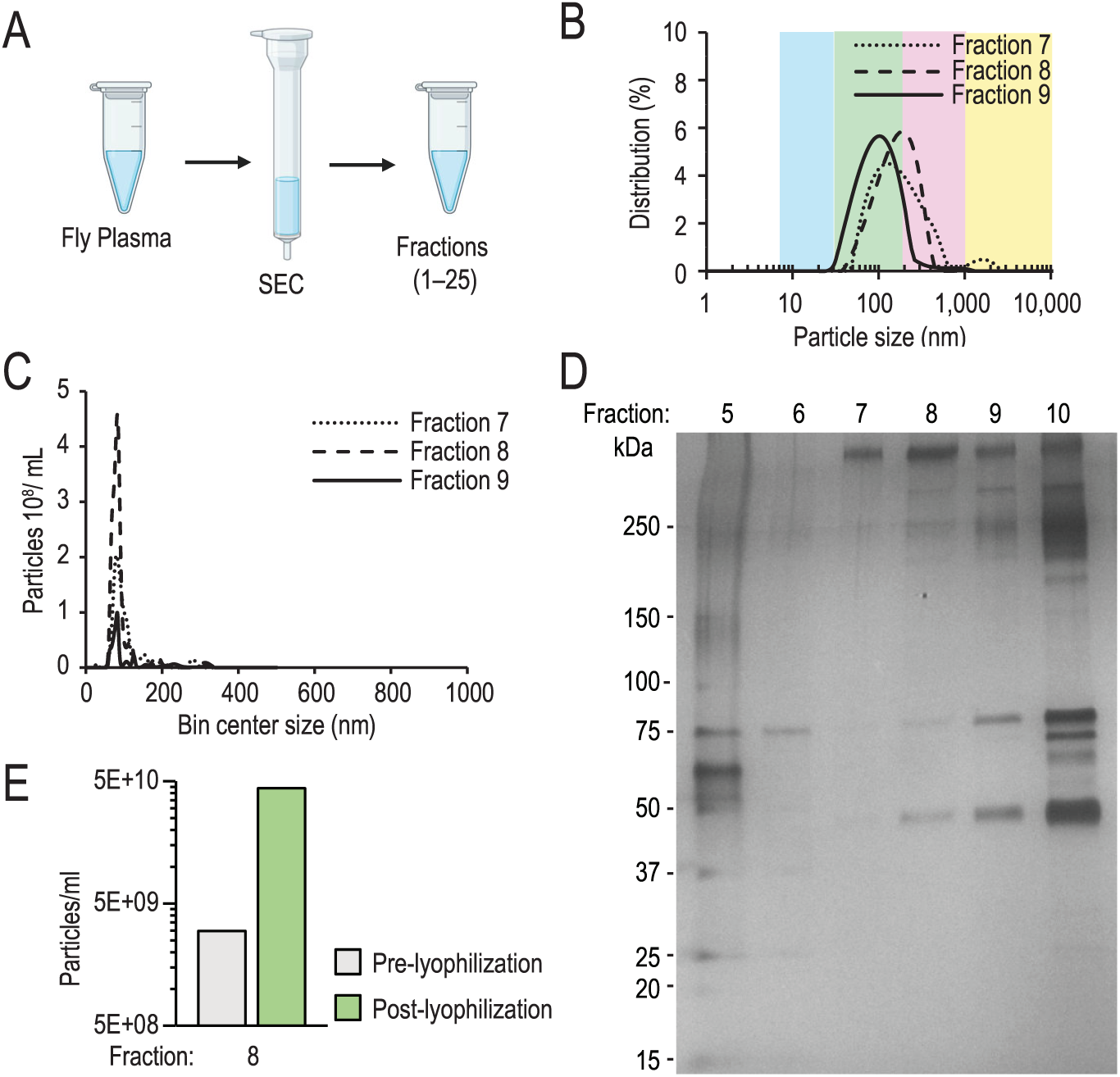
Size exclusion chromatography yields cleaner sEV fractions. (**A**) Schematic of *Drosophila* plasma fractionation using size exclusion chromatography (SEC). (**B**) DLS analysis of SEC fractions 7-9. Data was collected by DLS using three manual replicate measurements, with each run lasting 30 s. (**C**) Particle size distribution of fractions 7-9 measured by NTA. (**D**) Silver staining of SEC fractions 5-10. (**E**) Particle concentration measurements by NTA.

Next, we assessed the protein profiles of fractions 5–25 by silver staining. DLS detected large particles with a peak at 1000 nm in fraction 5, which were not observed in fractions 6–10 (Figure 5B and Supplementary Figure 1A). Consistent with this finding, the protein profile of fraction 5 was distinct from those of fractions 6–10 (Figure 5D). Small EV-sized particles were detected in fractions 6–13, whereas a significant portion of lipophorin-like particles smaller than 30 nm first appeared in fraction 10 and persisted through fraction 23 (Figure 5B, Supplementary Figure 1A). Presence of small EV-sized particles was negligible in fractions 16–21 (Supplementary Figure 1A). Correspondingly, the protein bands at ∼75 kDa and ∼250 kDa, which are thought to correspond to ApoLII and ApoLI, respectively, were prominently detected in fractions 11–22 (Supplementary Figure 1B). Notably, these ∼75 kDa and ∼250 kDa protein bands were also detected in fractions 7–9 although their intensity was much weaker than later fractions, such as fraction 17, the SP sEV fraction, and the UC sEV fraction (Figure 2D, 3E, 4E, and Supplementary Figure 1B). Because small EV-like particles were specifically detected in fractions 7–9, we examined whether their protein profiles differed from those of fraction 10 and later fractions. However, we failed to identify any protein bands that were specific to fractions 7–9 (Figure 5D, Supplementary Figure 1B).

Our data suggest that SEC separates small EVs from lipophorins more efficiently than SP or UC. However, because the small EVs were dispersed across multiple fractions, their concentration in each fraction was substantially lower than that obtained with the other methods (Figure 5E). Therefore, we investigated whether lyophilization could be used to concentrate SEC-isolated small EVs (Charoenviriyakul et al., 2018). Lyophilized fraction 8 was reconstituted in filtered PBS to one-tenth of the original volume, and the particle concentration and size distribution were compared before and after lyophilization. This approach successfully increased the particle concentration without altering the particle size distribution of fraction 8 (Figure 5E).

### SP and UC sEV fractions contain more circulating hemolymph proteins and apolipophorins compared to SEC sEV fraction

Thus far, we have shown that SP, UC, and SEC can enrich small EVs with varying degrees of purity and yield. To assess the ability of these techniques to separate circulating hemolymph proteins from small EVs and to enrich known small EV proteins, we preformed mass spectrometry (MS) on the sEV fractions. Our MS analysis detected 1151 proteins in the SP sEV fraction, 824 in the UC sEV fraction, and 773 proteins in the SEC sEV fraction, which was prepared by pooling fractions 7-9 (Supplementary Table 1). Pairwise correlation analysis indicated that the SP and UC sEV proteomes were the most similar to each other (Pearson correlation coefficient = 0.93), whereas the SEC sEV proteome shared fewer proteins with either the SP or UC sEV proteome (Pearson correlation coefficients = 0.37 for SEC vs. SP and 0.42 for SEC vs UC).

Given that hemolymph contains numerous circulating proteins, including the larval serum proteins (Lsp) Lsp1α, Lsp1β, and Lsp2, their removal is critical for selective enrichment of circulating small EVs. To approximate the abundance of proteins in each sEV fraction, we calculated the adjusted normalized spectral abundance factor (NSAF) values, spectral abundance normalized to protein sequence length, for each protein (see Methods). Notably, we found that the adjusted NSAF values of larval serum proteins were three- to five-fold lower in the SEC sEV fraction than in the SP and UC sEV fractions, suggesting that SEC separates circulating proteins more efficiently than SP and UC (Figure 6A).

**Figure 6.**
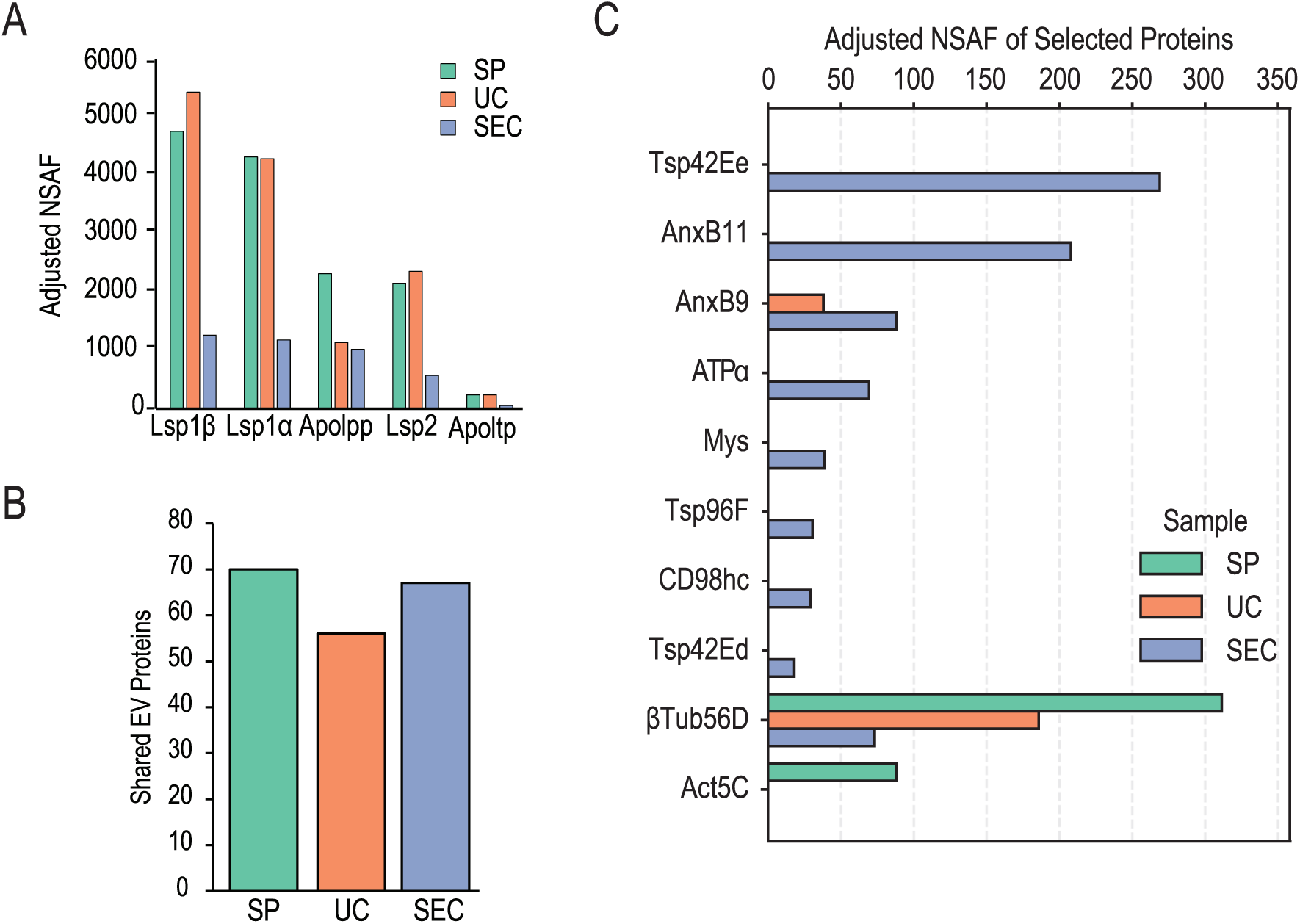
Comparison of the SP, UC, and SEC sEV proteomes. (**A**) Adjusted NSAF values for well-characterized circulating hemolymph proteins across all three techniques. (**B**) Total number of common proteins between sEV fractions and the curated top 100 EV protein dataset. (**C**) Adjusted NSAF values of select EV proteins in the SP, UC, and SEC sEV fractions.

Our findings indicate that lipophorins are also present in the sEV fractions. Apolpp and Apolipoprotein lipid transfer particle (Apoltp) are part of the *Drosophila* Apolipoprotein B (ApoB) protein family. The adjusted NSAF values for Apolpp in the UC and SEC sEV fractions were approximately two-fold lower than those in the SP sEV fraction (Figure 6A). We also detected lower levels of Apoltp in the SEC sEV fraction compared with the SP or UC sEV fractions (Figure 6A). These observations suggest that, while SEC removes lipophorins more efficiently than SP and UC, it does not eliminate them from the sEV fraction. The presence of Apolpp in the SEC sEV fraction is also consistent with our observation that the ∼75 and ∼250 kDa proteins are detected at low levels in fractions 7–9 (Figure 5D).

### SEC enriches small EV-associated proteins more efficiently than SP or UC

To assess whether previously reported small EV proteins are found in our sEV fractions, we compared our MS data with two public EV datasets: ExoCarta (Keerthikumar et al., 2016) and FunRich (Pathan et al., 2017), which is based on Vesiclepedia (Kalra et al., 2012). We compiled a list of proteins shared between the Top 100 EV protein lists curated by each database. Since ExoCarta and FunRich primarily curate mammalian EV proteins, we generated a list of the *Drosophila* orthologs for the shared EV proteins using the interspecies ortholog prediction tool DIOPT (Hu et al., 2011). A DIOPT score of at least 3 was applied when selecting *Drosophila* orthologs to remove low-confidence matches (see Methods). This generated a list of 189 *Drosophila* orthologs of the common EV proteins (Supplementary Table 2).

To remove proteins in our sEV fraction MS datasets that were likely detected randomly, we excluded low abundance proteins to generate filtered sEV fraction MS datasets (see Methods). We then compared our filtered sEV proteomes with the common EV protein *Drosophila* ortholog set and identified 70 proteins in the filtered SP sEV proteome, 56 proteins in the filtered UC sEV proteome, and 67 proteins in the filtered SEC sEV proteome that overlapped with the common EV protein *Drosophila* ortholog set (Figure 6B and Supplementary Figure 2). We also compared our filtered sEV proteomes with a published *Drosophila* small EV MS dataset (Linnemannstöns et al., 2022) and found that 309 shared proteins in the filtered SP sEV proteome, 250 shared proteins in the filtered UC sEV proteome, and 318 shared proteins in the filtered SEC sEV proteome (Supplementary Table 3).

*Drosophila* orthologs of several well-characterized small EV proteins were detected exclusively in the SEC sEV MS dataset, including Tetraspanin 42Ee (Tsp42Ee), Tetraspanin 42Ed (Tsp42Ed), Tetraspanin 96F (Tsp96F), Myospheroid (Mys), CD98 heavy chain (CD98hc), Na pump α subunit (Atpα), and Annexin B11 (AnxB11). Tsp42Ee and Tsp42Ed are the *Drosophila* orthologs of mammalian CD63, a well-established exosome marker (Conde-Vancells et al., 2008; Garcia et al., 2015; Gurung et al., 2021; Palmulli et al., 2024). Tsp96F is the *Drosophila* ortholog of mammalian CD81, another well-established exosome marker (Conde-Vancells et al., 2008; Garcia et al., 2015; Gurung et al., 2021). Additionally, ATP1A1, the mammalian ortholog to *Drosophila* Atpα, has been proposed as an exosomal marker candidate across multiple mammalian cell lines (Kugeratski et al., 2021). Interestingly, we also identified Mys (the *Drosophila* integrin β1 homolog) and one of its binding partners, CD98hc, in the SEC sEV dataset (Figure 6C). Both proteins have been previously identified in mammalian exosomes (Kugeratski et al., 2021). Some human annexins play critical roles in membrane dynamics and EV biogenesis (Bischoff et al., 2013; Gradilla et al., 2014). Interestingly, two *Drosophila* Annexins, AnxB11 and AnxB9, were also enriched in the SEC sEV proteome compared to the SP and UC sEV proteomes. These data demonstrate that the SEC sEV fraction contains more high-confidence EV proteins than the SP or UC sEV fractions. Note that not all proteins shared between our sEV datasets and the common EV protein *Drosophila* ortholog set were more abundant in the SEC sEV fraction compared to the SP sEV and UC sEV fractions. For example, cytoskeletal proteins βTub56D (the *Drosophila* ortholog of TUBB4B) and Act5C (the *Drosophila* ortholog of ACTB), were more abundant in the SP sEV fraction than the UC sEV or SEC sEV fractions (Figure 6C).

### Further characterization of the proteins unique to the SEC sEV fraction revealed multiple exosome-related features

Given that well-established exosomal markers, such as tetraspanins and integrin subunits, were exclusively identified in the SEC sEV proteome (Figure 6C), we further characterized proteins unique to the SEC sEV proteome using the analysis tools available in the Search Tool for the Retrieval of Interacting Genes/Proteins (STRING) database (https://string-db.org/) (Szklarczyk et al., 2023). After identifying proteins unique to the SEC sEV proteome (Supplementary Table 4), we determined their human orthologs using DIOPT because human protein annotations and interaction networks are more comprehensively documented. We excluded proteins with low DIOPT scores (< 3). When multiple human orthologs were predicted, we selected the ortholog with the highest DIOPT score. If all predicted human orthologs shared the same DIOPT score, we randomly selected a single representative ortholog. Ultimately, we identified human orthologs for 108 of the 140 *Drosophila* proteins found exclusively in the SEC sEV proteome (Supplementary Table 5). These 108 proteins were enriched in several biological processes, including transport, localization, vesicle-mediated transport, organic substance transport, and viral life cycle (Figure 7A and Supplementary Table 6). Cellular component gene ontology (GO) enrichment analysis revealed that these proteins associated with the lysosomal membrane, extracellular exosome, cytoplasmic vesicle, and late endosome membrane (Figure 7B and Supplementary Table 7). Because exosomes are generated within MVBs, which are specialized late endosomes that either fuse with lysosomes for exosome degradation or with the plasma membrane to release exosomes extracellularly, the enrichment of late endosomal, lysosomal, and vesicle-associated GO terms strongly supports the conclusion that the SEC sEV fraction is enriched for proteins associated with exosome biogenesis and secretion.

**Figure 7.**
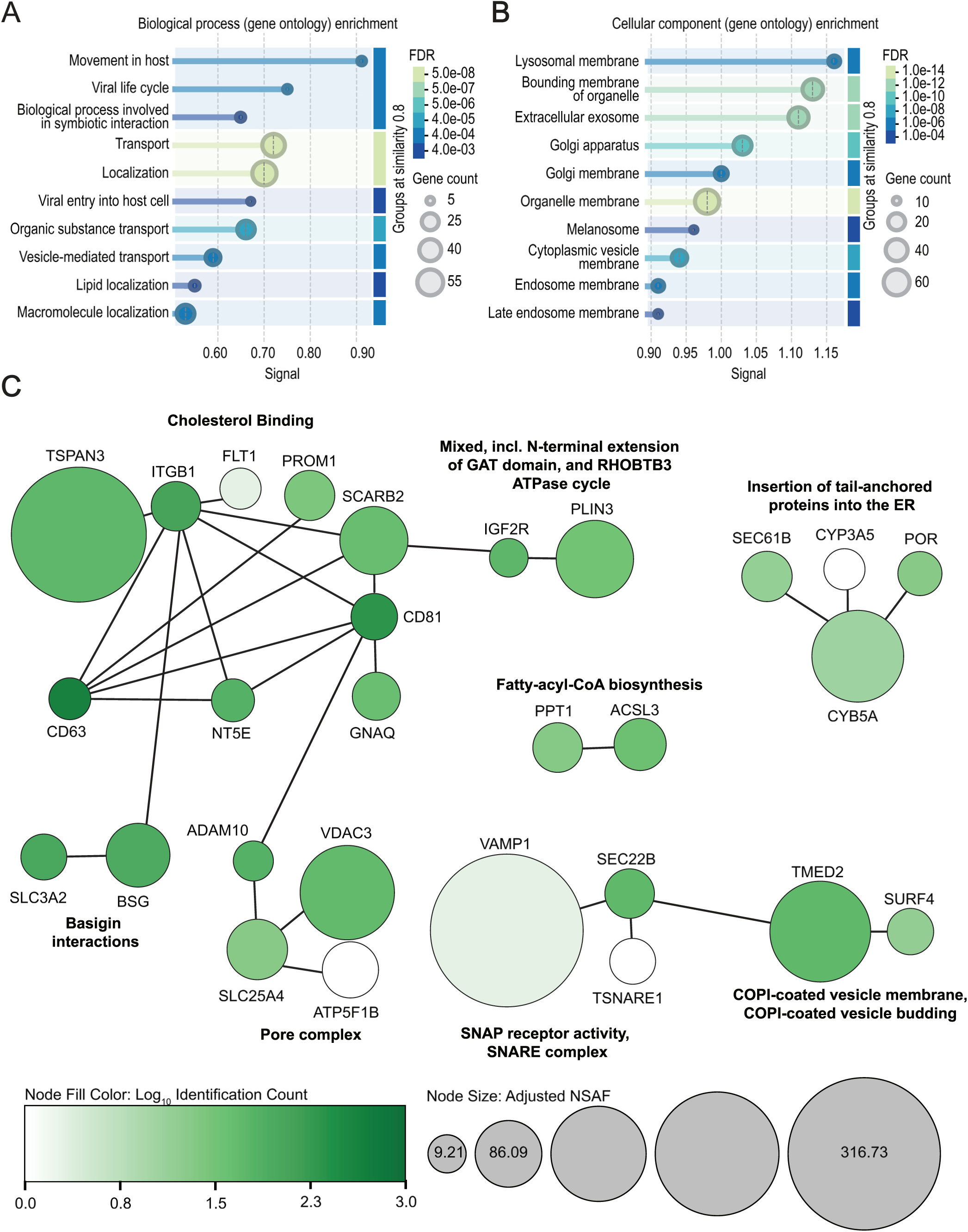
SEC-exclusive proteins contain protein clusters involved in small EV biogenesis and function. (**A-B**) Gene ontology enrichment analysis of SEC-exclusive proteins for biological process (**A**) or cellular component (**B**). (**C**) Clustering identified 8 functional groups. Biological terms associated with each cluster were generated by STRING-DB. Solid edges represent established interactions. Node color correlates to the number of ExoCarta-curated experiments in which the protein was identified (see Methods). Node size is scaled according to adjusted NSAF value in the SEC sEV fraction.

To gain further insight into these 108 SEC sEV exclusive proteins, we retrieved known high-confidence interactions among them and identified eight distinct clusters using Markov clustering algorithm (inflation parameter = 1.8). We colored each node according to the number of ExoCarta-curated experiments in which the corresponding protein was documented (identification count) and scaled its size based on the adjusted NSAF value (see Methods) (Figure 7C). Interestingly, the high-confidence interaction network included several well-established EV proteins, including CD63, CD81, integrin subunit beta 1 (ITGB1), transmembrane emp24 domain-containing protein (TMED), and a disintegrin and metalloproteinase domain-containing protein 10 (ADAM10), all of which exhibited high identification counts (Figure 7C). We then performed functional enrichment analysis for each cluster and highlighted the most significantly enriched terms (Figure 7C). Among the eight clusters, the largest cluster was enriched for proteins with ‘cholesterol binding’ activity (Figure 7C). Interestingly, intracellular cholesterol levels have been shown to regulate exosome secretion (Möbius et al., 2003). Moreover, CD63 has recently been reported to regulate cholesterol trafficking by sorting cholesterol into nascent exosomes before their release (Palmulli et al., 2024). Other notable functional clusters include ‘SNARE complex’, ‘insertion of tail-anchored proteins into the ER membrane’, ‘pore complex’, and ‘COPI-coated vesicle membrane proteins’ (Figure 7C). These functional categories are all associated with biological membranes, suggesting that the SEC sEV proteome is enriched for membrane-associated proteins, including proteins commonly found in exosomes.

Taken together, our analysis of the proteins unique to the SEC sEV fraction uncovered several features associated with small EVs. These results further support the conclusion that the SEC sEV fraction contains small EV-related proteins at higher abundance than the SP and UC sEV fractions.

## Discussion

Due to their simplicity and experimental tractability, tissue culture systems have served as the primary platform for small EV research; however, these systems often fail to faithfully recapitulate *in vivo* small EV biology (Böröczky et al., 2023; Couch et al., 2021; J. C. Lee et al., 2024; Van Niel et al., 2022). Consequently, there is a growing need for simple yet robust, genetically tractable *in vivo* model systems to study small EVs (Böröczky et al., 2023; Couch et al., 2021; J. C. Lee et al., 2024; Van Niel et al., 2022). Given its simplicity and advanced genetic tools, *Drosophila* is a powerful animal model for investigating the fundamental biology of small EVs in normal physiology and disease (Ajibefun et al., 2026; Ogienko et al., 2022). Although *Drosophila* has been used to study small EVs in the context of cell biology (Blanchette et al., 2022; Camelo et al., 2022; Gross et al., 2012; Hurbain et al., 2022; Koles et al., 2012; Lee et al., 2026; Linnemannstöns et al., 2022), the isolation of small EVs for biochemical and functional analyses remains challenging. Here, we evaluated whether conventional small EV enrichment methods, including SP, UC, and SEC, can be directly applied to isolate small EVs from *Drosophila* hemolymph. We provide a comprehensive comparison of the performance and limitations of three widely used enrichment methods for isolating small EVs from *Drosophila* hemolymph.

### Our DLS analyses reveal four distinct particle populations in *Drosophila* hemolymph

Our observations suggest that the smallest particles, ranging from ∼10 to ∼30 nm, represent lipophorins, and the particles ranging from 50 to 200 nm represent small EVs (Figures 1D, 4, and 5). DLS also detected particles larger than 2,000 nm, likely representing cell debris and small hemocytes, as well as particles ranging from 200 to 600 nm, whose size distribution and sedimentation properties are reminiscent of the large EVs described in mammals. Particles larger than ∼300 nm can be efficiently removed by a series of low-speed centrifugation steps (Figure 1D). Removing these large particles prior to small EV enrichment is crucial for minimizing contamination by larger particles, as they can co-sediment with small EVs during SP and UC. Notably, preventing hemolymph coagulation and melanization is also critical for obtaining a cleaner small EV preparation (Figure 1C). Thus, we prepared *Drosophila* “plasma” by adding PTU and EDTA, followed by a series of low-speed centrifugation steps (Figure 1B), before proceeding with small EV enrichment procedures.

### Conventional small EV preparation procedures enrich small EVs to varying degrees

Our DLS and NTA analyses of plasma revealed a particle population with a peak diameter of approximately 100 nm. Their size distribution and sedimentation properties were reminiscent of small EVs found in mammals (Kalluri & LeBleu, 2020). These particles were present in the small EV fractions prepared by SP, UC, and SEC (Figures 2B, 2C, 3B, 3C, 5B, and 5C). Notably, cryo-electron microscopy of the UC sEV fraction revealed vesicles ranging from approximately 70 to 200 nm in diameter (Figure 3D). Our DLS and silver staining analyses showed that SEC yielded the cleanest small EV fractions. Nevertheless, we failed to detect protein bands that appeared to be specific to small EVs in the SEC sEV fractions (fractions 7–9) or the SP sEV fractions (Figure 2D and 5D). While two distinct protein bands were observed in the UC sEV fractions (Figure 3E), none of the top 30 enriched proteins in the UC sEV fraction were recognized as a well-established small EV protein (Supplementary Table 1). NTA analysis showed that the total number of particles recovered by SEC was comparable to that recovered by UC, although both yielded fewer particles than SP. However, the protein content of the SEC sEV fraction was consistently below the detection limit of conventional protein quantification assays. The low protein abundance of the SEC sEV fraction therefore poses a significant challenge for characterizing them. Although lyophilization successfully increased particle concentration (Figure 5E), it failed to raise protein concentrations to measurable levels under our experimental conditions. Scaling up the isolation procedure or employing more sensitive protein detection methods may help overcome this limitation.

### Established exosome markers are enriched in the SEC small EV fraction compared with the SP and UC sEV fractions

Our mass spectrometry analysis revealed that the SEC sEV fraction was enriched for small EV-associated proteins, including *Drosophila* orthologs for CD63, CD81, ATPA1, and Annexin B, which were not detected in the SP or UC sEV fractions (Figure 6C). GO enrichment analysis identified several biological process and cellular component terms associated with small EVs, including ‘vesicle-mediated transport’, ‘extracellular exosome’, ‘cytoplasmic vesicle membrane’, and ‘endosome membrane’ (Figure 7A and 7B). STRING analysis of proteins uniquely detected in the SEC sEV fraction further identified functional clusters previously reported to be involved in exosome biogenesis, such as ‘SNARE complex components’, ‘COPI vesicle membrane’, ‘pore complexes’, and ‘cholesterol binding’ (Figure 7C). Collectively, these findings demonstrate that although SEC yields less total protein, it reliably enriches small EVs from *Drosophila* hemolymph when compared with SP and UC. In addition, SEC was more time-efficient and produced more reproducible results than UC or SP.

### None of the techniques completely separate lipophorins and circulating hemolymph proteins from small EVs

Our genetic knockdown experiments indicate that the protein bands detected at ∼75 and ∼250 kDa represent the two major protein components of lipophorins, ApoLII and ApoLI, respectively (Figure 4A and E). The SP procedure failed to reduce the intensity of these protein bands, as well as those of other major circulating proteins present in the hemolymph (Figure 2D). Consistently, DLS detected lipophorins in the SP sEV fraction (Figure 2B). Although the UC procedure substantially removed lipophorins and other major circulating proteins, the intensity of the ∼75 and ∼250 kDa protein bands were not significantly reduced in the UC sEV fraction (Figure 3B and E). In contrast, the intensity of these bands, along with other major protein bands, was markedly reduced in SEC fractions 7–9, which were enriched for small EV-sized particles, compared with the SP and UC sEV fractions (Figure 5D). Consistent with these observations, lipophorins (<30 nm) were not detected in SEC fractions 7–9 by DLS (Figure 5B and C). Interestingly, the presence of lipophorins detected by DLS closely correlated with the intensity of the ∼75 and ∼250 kDa protein bands (Supplementary Figure 1A and B). For example, DLS detected abundant lipoprotein-like particles in later fractions, such as fraction 17, where the intensity of the ∼75 and ∼250 kDa protein bands were the highest (Supplementary Figure 1A and B). These observations are consistent with our mass spectrometry analysis. The adjusted NSAF values of apolipoproteins (Apolpp and Apoltp) and larval serum proteins were lower in the SEC sEV fraction than in the SP and UC sEV fractions (Figure 6A and Supplementary Table 1).

Together, these findings indicate that SEC is more effective than SP and UC at separating lipoproteins and major circulating proteins from small EVs. Nevertheless, none of the three techniques completely removed Apolpp from the final sEV fractions. This persistent presence of Apolpp may reflect the high abundance of lipophorins in *Drosophila* hemolymph. Alternatively, the physical properties of *Drosophila* lipophorins may differ from those of mammalian lipoproteins, making them more difficult to separate using conventional small EV isolation procedures. Given that NTA has limited sensitivity for detecting lipoprotein-sized particles, combining DLS and silver staining with NTA should provide a more comprehensive characterization of small EV preparations for downstream cargo discovery and functional assays.

### We present a proteomic characterization of small EV fractions isolated from larval *Drosophila* hemolymph

We report high-quality proteomes of sEV fractions isolated by SP, UC, and SEC, serving as reference proteomes for future sEV research in *Drosophila* (Supplementary Table 8). Our study demonstrates that the sEV fraction prepared by SEC contains *Drosophila* orthologs of well-established sEV marker proteins (Figure 6C and 7C; Supplementary Figure 2; Supplementary Table 5). Consequently, proteins identified exclusively in the SEC sEV fraction may include novel *Drosophila* sEV proteins. We anticipate that our proteomics datasets—particularly the SEC sEV-exclusive protein list—will serve as a valuable resource for discovering new small EV proteins and increasing the rigor of future small EV studies in *Drosophila*.

### Perspectives on the isolation and analysis of small EVs from *Drosophila* hemolymph

*In vivo* studies provide vital insights into the physiological functions of EVs (Welsh et al., 2024). Such studies typically involve either examining endogenous EVs or reintroducing enriched EV fractions into an organism. In both cases, the quality of EV preparations and careful characterization of the isolated EV fractions are critical for drawing reliable conclusions. Our study provides practical guidance on how to use these small EV isolation methods to achieve optimal enrichment and how to thoroughly characterize the resulting small EV fractions. Given that hemolymph is composed of particles varying in size, such as lipoproteins, small EVs, large EVs, and hemocytes (Figure 1D), we emphasize the use of particle analysis tools detecting a wide size range, such as DLS, for rigorous characterization of the prepared small EV fractions. Precise measurement of particle concentrations using assays such as NTA is also crucial. Furthermore, analyzing the protein profile or content using silver staining and mass spectrometry should provide a reliable assessment on how much circulating proteins and apolipoproteins are co-enriched with small EVs. Although we sought to establish clear guidelines for applying small EV enrichment methods to *Drosophila* hemolymph, our findings of the three commonly used small EV isolation methods were somewhat disappointing. None of these methods yielded small EV fractions of sufficient purity and quantity for direct biochemical characterization of their cargo or for transplantation experiments to elucidate their functions. To address the limitations of these small EV isolation methods, further optimization and methodological refinement may be necessary. Additionally, combining these methods may help overcome the limitations of each individual method. Ultimately, our findings highlight the need for a novel strategy that overcomes the limitations of current small EV isolation methods. Such a strategy will advance the use of *Drosophila* as a genetically tractable *in vivo* model for rigorous small EV research. Beyond *Drosophila*, the knowledge gained from this study provides a framework for the isolation and characterization of circulating small EVs in other insects, thereby facilitating future EV research across diverse insect species.

## Materials and Methods

### *Drosophila* stocks and genetics

Stable lines and fly crosses used for small EV isolation were maintained in bottles with standard cornmeal-agar medium and kept at room temperature throughout development. Stocks obtained from the Bloomington *Drosophila* Stock Center (BDSC) are as follows: *UAS-apolpp-RNAi^HM05157^* (BDSC, 28946), *w^1118^* (BDSC, 3605), *Cg-GAL4* (BDSC, 7011).

### Hemolymph collection and preparation

Hemolymph collection buffer composed of 2 mM EDTA (Sigma-Aldrich, ED2SS-500G) and 1 ug/ml phenylthiourea (Fisher Scientific, AAL0669009) in 1X PBS (Fisher Scientific, 10-010-031) was made and filtered through sterile 0.22 µm PES filters (VWR, 76479-044). Filtered hemolymph collection buffer was analyzed via ZetaView nanoparticle tracking analyzer (Particle Metrix) for any particle contamination.

For each small EV isolation, hemolymph was collected from 200 wandering third instar larvae. Larvae were separated from food using a 0.5 M sucrose solution. Larvae were transferred to a new 50 mL conical containing 40 mL 0.5 M sucrose solution and rinsed to remove food debris. Larvae were transferred to a new 50 mL conical containing dH_2_O and rinsed 3 times to remove the sucrose solution. Larvae were dried on a KimWipe and transferred into a 100 µl droplet of hemolymph collection buffer on Parafilm. Forceps were used to carefully puncture the larval cuticle, ensuring minimal damage to internal organs. Hemolymph from batches of larvae with visible organ damage were not used. After bleeding, hemolymph collection buffer was retrieved, transferred into a microcentrifuge tube, and kept on ice. Hemolymph was subjected to 3 rounds of centrifugation at 4 °C, transferring supernatant between spins: 500 x *g* 10 min, 2,000 x *g* 10 min, 10,000 x *g* 30 min. Final supernatant, termed “plasma” was used for all small EV isolation experiments. Collection of hemolymph from 200 larvae yielded approximately 150 µl of plasma.

### Ultracentrifugation

The entire plasma sample was ultracentrifuged in a 0.2 mL open-top thickwall polycarbonate tube (Beckman Coulter, 343775) using a TLA-100 fixed-angle rotor (Beckman Coulter, 343840) for 70 minutes at 100,000 x *g*, 4 °C. Approximately 150 µl of supernatant was collected and saved, termed “supernatant 1 (SN1)”. The pellet was resuspended in 150 µl of filtered PBS and ultracentrifuged again. Approximately 150 µl of supernatant was collected and saved, termed “supernatant 2 (SN2)”. The UC small EV pellet was resuspended in 50 µl filtered PBS.

### Solvent precipitation

50 µl of XENO-EVI Pre-Buffer (XENOHELIX, 9366-EVI) was added to 100 µl of *Drosophila* plasma, briefly vortexed to mix, and centrifuged 14,000 x *g* for 15 minutes at 4 °C. The supernatant was transferred to a new microcentrifuge tube where 40 µl of XENO-EVI buffer was added. Sample was briefly vortexed and incubated for 10 minutes at room temperature. After incubation, the sample was centrifuged 12,300 x *g* for 10 minutes at 25 °C. The supernatant was discarded and the pellet, termed “SP sEV” was resuspended in 50 µl filtered PBS.

### Size exclusion chromatography

During *Drosophila* plasma preparation, a 35 nm qEVsingle column (IZON, ICS-35) was mounted in an upright position over a waste collection beaker. The bottom, then top cap was removed, and buffer was allowed to drain. The qEV column was washed twice with 3 mL of PBS which had been filtered and equilibrated to room temperature. After the second wash, the column was positioned over an open microcentrifuge tube. A total of 150 µl *Drosophila* plasma was loaded into the column and displaced buffer was collected. SEC fractions were eluted by adding 170 µl of PBS, filtered and equilibrated to room temperature, to the column and collecting the corresponding fraction. For small EV isolations, 15 fractions were collected. For whole hemolymph fractionation, 25 fractions were collected. All fractions were analyzed by DLS immediately and snap frozen in liquid nitrogen.

### Particle size analysis

All small EV fractions were analyzed by dynamic light scattering (Anton Paar Litesizer 100). The Omega cuvette was rinsed 3 times with MilliQ water. Samples were measured using a particle size series with three manual replicate runs of 30 seconds each. Nanoparticle tracking analysis was conducted using a ZetaView particle analyzer (Particle Metrix). Instrument sensitivity was set to 80 and shutter to 150. All samples were diluted in filtered PBS as follows: UC sEV, 1:100; SP sEV, 1:100; SEC, 1:50.

### Electron microscopy

Purified small EV samples were recovered after DLS analysis. Five microliters of small EV samples were applied to glow-discharged CFLAT 2/2 holey-carbon grids (Protochips), blotting four times, then plunging into liquid ethane using a Vitrobot (ThermoFisher). Exosome samples were imaged at 67,000X magnification using a Glacios (ThermoFisher) equipped with a K2 summit direct detection camera (Gatan Inc.) as well as a Quantum Gatan Imaging Filter energy filter (Gatan Inc).

### Immunoblot analysis

Small EV samples were lysed in buffer containing 50 mM Tris-HCl, pH 8, 0.1% SDS, 0.5% sodium deoxycholate, 1% NP-40, 150 mM NaCl and Halt Protease and Phosphatase Inhibitor Cocktail (ThermoFisher Scientific, 78442). Protein concentration was measured by ThermoFisher Scientific Pierce BCA Protein Assay Kit. Samples were prepared in 4X Laemmli sample buffer (Bio-Rad 16710747) supplemented with 355 mM 2-mercaptoethanol and boiled for 5 minutes at 95 °C. 2 µg protein samples were separated on 4-15% Mini-PROTEAN TGX Precast Protein Gels (Bio-Rad, 4561086) and transferred to PVDF membranes using Trans-Blot Turbo PVDF Transfer packs (Bio-Rad, 1704156) in the Trans-Blot Turbo Transfer System (Bio-Rad, 1704150). Membranes were blocked in 5% non-fat dry milk in 1X TBS-T for 1 hour at room temperature and then incubated in rabbit-ApoLII (1:500, gift from Dr. Akhila Rajan) in blocking buffer overnight at 4 °C. Membranes were rinsed 3 times in 1X TBS-T and incubated in IRDye 800CW goat anti-rabbit IgG antibody (1:2000, LICOR Biosciences, 926-32211) for 45 minutes at room temperature. Membranes were rinsed 3 times in 1X TBS-T and imaged using a LI-COR CLx Odyssey western blot scanner.

### Mass spectrometry

To prepare samples for mass spectrometry (MS) analysis, we directly lysed, reduced, digested, and cleaned up the lyophilized small EV fractions which had been stored at -80 °C using the EasyPep Mini MS sample prep kit (ThermoFisher Scientific, A40006). We dried the prepared peptides overnight in a SpeedVac and stored them at - 80 °C until MS analysis. LC-MS/MS was performed as previously described (K.-T. Lee et al., 2024).

### Mass spectrometry data processing

MS/MS spectra were queried and filtered as previously described (K.-T. Lee et al., 2024). After the results were processed using the Trans-Proteomic Pipeline suite of tools (Keller et al., 2002; Nesvizhskii et al., 2003), the resulting pepXML and protXML files were used as input for Abacus (Fermin et al., 2011), a computational tool that extracts spectral counts from MS/MS data to accurately adjust them to account for peptides shared among multiple proteins. The resulting spectral counts were normalized to generate adjusted normalized spectral abundance factors (adjusted NSAFs) (Zhang et al., 2010) (Supplementary Table 8). Before analyzing the proteomes of the sEV fractions, we removed proteins with adjusted NSAF values of <2 to filter out proteins that were likely detected randomly.

Human orthologs for *Drosophila* proteins were annotated using the DRSC Integrative Ortholog Prediction Tool (DIOPT) (Hu et al., 2011). To remove low-confidence ortholog predictions, we used a DIOPT score of ≤ 2 as the cutoff for both the common EV protein *Drosophila* ortholog set and the human orthologs exclusive to the SEC sEV dataset.

To quantify how frequently each SEC sEV-exclusive protein was identified across different experiments in the ExoCarta database, we downloaded the entire ExoCarta protein dataset and filtered out non-*Homo sapiens* entries. We then determined the total number of curated experiments in ExoCarta that detected each of the 108 SEC sEV-exclusive human orthologs, which we termed the ‘identification count.’ Protein–protein interactions and functional clusters among the 108 SEC sEV-exclusive human orthologs were identified using the STRING database (Szklarczyk et al., 2023). The resulting network was visualized in Cytoscape (Shannon et al., 2003). The log_10_ identification count was mapped to node color, and adjusted NSAF values were mapped to edge thickness.

### Schematics

Figure 1A was made using Adobe Illustrator. All other schematics were made using BioRender.

## Acknowledgements

We thank Joel Quispe for assistance with cryo-EM, Dr. Akhila Rajan for providing the anti-ApoLII antibody, and Jimmy Eng for help with processing raw mass spectrometry data. This work was supported by Kuni Foundation Discovery Grants for Cancer Research to Y.V.K.

## Author Contributions

A.G. and Y.V.K. designed experiments, analyzed data, and wrote the manuscript. C.G.V. analyzed data and wrote the manuscript. A.G. performed experiments. Y.V.K. acquired funding and supervised the project. S.W.Y reviewed and edited the manuscript.

## Supplementary Figure Legends

**Supplementary Figure 1.**
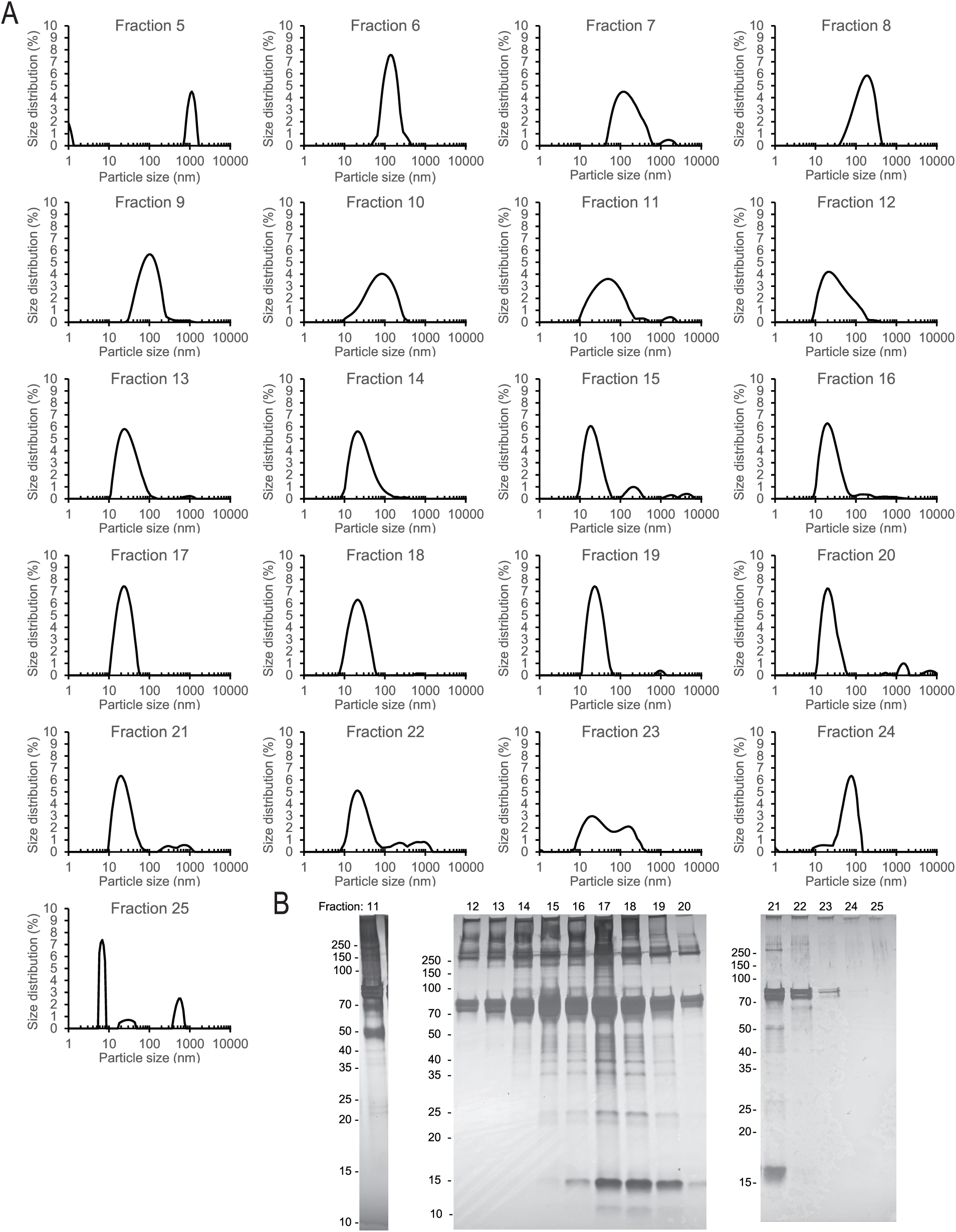
Particle and protein analysis of SEC fractions obtained from hemolymph. (**A**) Particle size distributions for all 25 SEC hemolymph fractions. Data was collected by DLS using three manual replicate measurements, with each run lasting 30 s. (**B**) Silver staining of SEC hemolymph fractions.

**Supplementary Figure 2.**
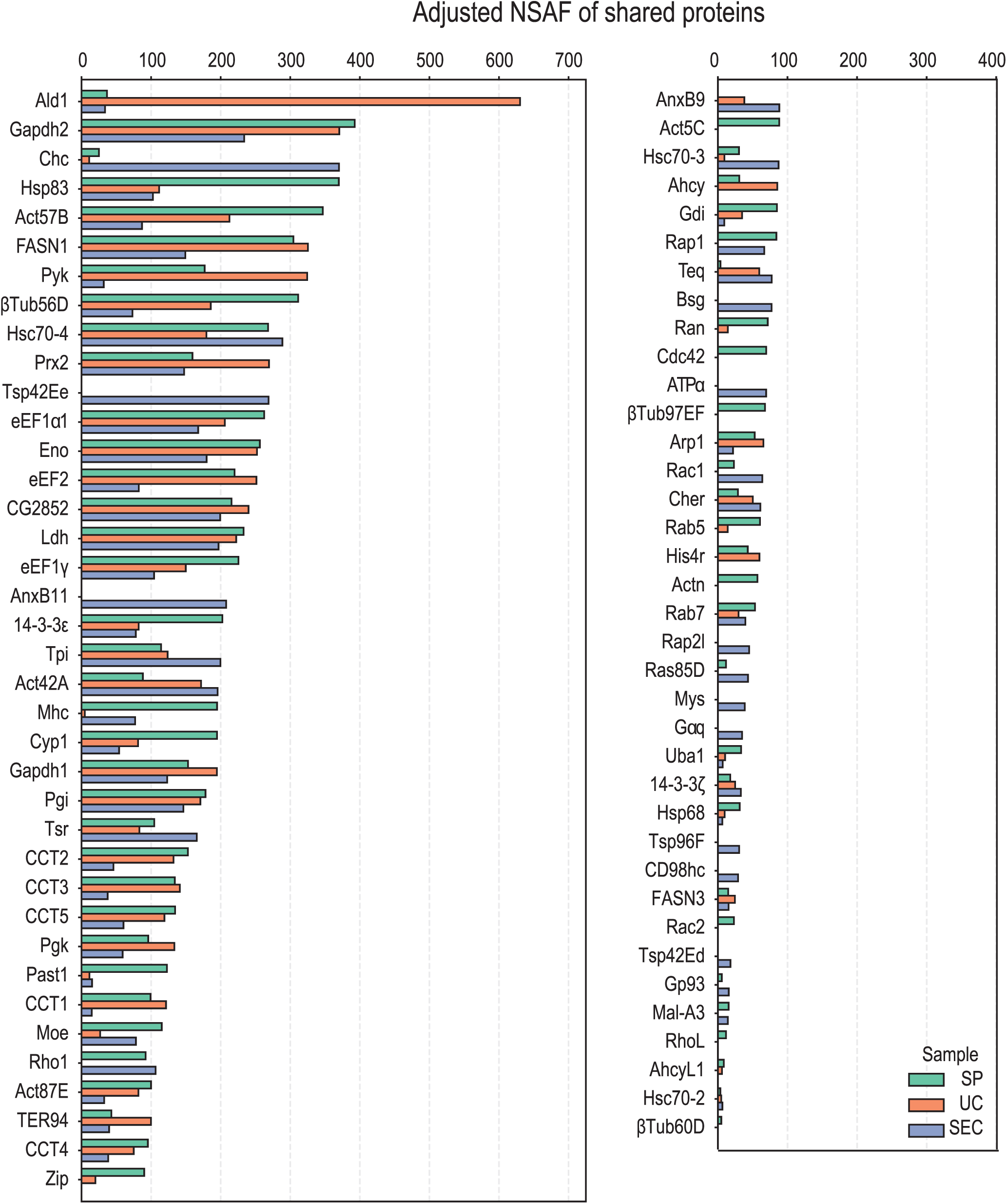
Common proteins between the common EV protein *Drosophila* ortholog set and the filtered sEV fraction MS datasets. Adjusted NSAF values for each protein in each sEV fraction are plotted on the x-axis to indicate protein abundance.

## Supplementary Table Legends

**Supplementary Table 1. Adjusted NSAF values for proteins detected by mass spectrometry in the SP, UC, and SEC sEV fractions.**

**Supplementary Table 2. The common EV protein *Drosophila* ortholog set.** First, we identified common proteins between the top 100 EV protein lists curated by the ExoCarta and FunRich databases. Then, we used the DIOPT tool to find *Drosophila* orthologs for the shared proteins. We used a DIOPT score of ≤ 2 as the cutoff to remove low-confidence ortholog predictions,

**Supplementary Table 3. Shared proteins between the filtered SP, UC, and SEC sEV fraction proteomes and and the *Drosophila* small EV dataset reported by Linnemannstöns et al. (ref).** The total number of shared proteins between each sEV fraction and the published *Drosophila* small EV dataset is shown in the “Summary” sheet. The ‘SP’ sheet contains the names of proteins shared between the published Drosophila small EV dataset and the SP sEV fraction, along with the adjusted NSAF value for each shared protein in the SP sEV fraction. The ‘UC’ sheet contains the names of proteins shared between the published Drosophila small EV dataset and the UC sEV fraction, along with the adjusted NSAF value for each shared protein in the UC sEV fraction. The ‘SEC’ sheet contains the names of proteins shared between the published Drosophila small EV dataset and the SEC sEV fraction, along with the adjusted NSAF value for each shared protein in the SEC sEV fraction.

**Supplementary Table 4. Proteins exclusively found in the SEC sEV fraction.** Filtered sEV fraction proteomes were compared and proteins with a SEC sEV adjusted NSAF score of ≥ 2 and SP and UC sEV adjusted NSAF score of = 0 were retained.

**Supplementary Table 5. Human orthologs to the 140 SEC sEV exclusive proteins.** Predicted human orthologs for *Drosophila* SEC sEV exclusive proteins were generated using the DIOPT ortholog prediction tool (Hu et al., 2011). Human orthologs with a DIOPT score < 3 were removed. For each *Drosophila* protein, the human ortholog with the highest DIOPT score is specified in column B, “Human ortholog 1”. For *Drosophila* proteins with multiple predicted human orthologs with tied DIOPT scores, one ortholog was chosen at random.

**Supplementary Table 6. Gene Ontology biological processes enrichment terms for proteins exclusively found in the SEC sEV fraction.** The analysis was performed using the STRING database.

**Supplementary Table 7. Gene Ontology cellular component enrichment terms for proteins exclusively found in the SEC sEV fraction.** The analysis was performed using the STRING database.

**Supplementary Table 8. Unprocessed sEV fraction mass spectrometry data.** The “UniProt” sheet contains the unprocessed sEV fraction mass spectrometry data. In the “UniProt” sheet, each unique protein is identified by the UniProt KB accession identifier (PROTID). To annotate the *Drosophila* protein name, the PROTID was used to identify the corresponding FlyBase FBpp number in the “FlyBase” sheet. The FBpp number was used to identify both the CG number and isoform-specific gene name of each *Drosophila* protein, in columns “CG” and “gene”, respectively.

